# Characterizing the Binding Mechanism of Sparsentan to the Type-2 Angiotensin II Receptor Using AutoDock Vina

**DOI:** 10.1101/2024.03.22.586129

**Authors:** Peter X. Geng

## Abstract

The angiotensin II type-2 receptor (AT2R) is known to have a significant impact on cardiovascular physiology, and the elucidation of its drug interactions is crucial for advancing therapeutic interventions. Sparsentan is an antagonist of angiotensin II, which has received FDA approval due to its demonstrated efficacy in ameliorating proteinuria. While sparsentan exhibits a preferential affinity for angiotensin II type-1 receptor (AT1R) and may primarily target it, DrugBank identifies it as a drug that targets AT2R. Present databases lack detailed structural data on the interaction mechanism between sparsentan and AT2R, leaving no definitive evidence to confirm or refute an interaction with AT2R. This study aimed to explore the potential for an interaction between sparsentan and AT2R. Molecular docking simulations were conducted using AutoDock Vina to predict the binding conformation. The docking parameters were meticulously optimized to ensure the accuracy and reliability of the simulation results. Subsequently, the most stable sparsentan-AT2R complex was determined. We identified four primary types of interactions at the binding site containing hydrogen bonds, hydrophobic interactions, π-cation interactions, and π-stacking, that are likely to contribute to the affinity of sparsentan for AT2R. Our study indicates that sparsentan may potentially interact with AT2R, our results may serve as a valuable reference for elucidating the mechanism of sparsentan action and for guiding the design of AT2R-targeted drugs.

## Introduction

### Angiotensin II Type-2 Receptor (AT2R)

The renin-angiotensin system (RAS) is essential in maintaining cardiovascular stability and balance. The primary active component of this system is angiotensin II (Ang II) that exerting its effects mainly through interaction with two primary types of G-protein coupled receptor (GPCR): the angiotensin II type-1 receptor (AT1R) and the AT2R^1^. The genes encoding the AT2R are located on human chromosome X^2^. AT1R and AT2R are members of the seven-transmembrane domain receptor superfamily, with a 34% similarity in their nucleotide sequences^3^. AT2R exhibits high levels of expression in fetal tissue, but this expression markedly decreases and becomes confined to select organs, such as those within the cardiovascular system. The AT2R is also re-expressed in the adult animal playing a role for tissue remodeling, growth and development^4^.

AT2R functions as a modulator of blood pressure and electrolyte balance, exhibiting effects often opposite to those mediated by the AT1R. It promotes vasodilation and apoptosis, which are thought to contribute to its cardioprotective and antihypertrophic effects^5^. The receptor activates a number of intracellular signaling pathways involving phosphatases such as mitogen-activated protein kinase (MAPK) phosphatase-1, and can modulate ion transport across membranes, influencing cellular proliferation and differentiation^6^. Dysregulation of AT2R expression or function has been implicated in various diseases. For example, altered AT2R signaling is associated with hypertension, heart failure, and renal diseases. Additionally, AT2R’s presence in the brain suggests it may play a role in neurological conditions, which is an area of ongoing research^7^.

Recent studies focus on elucidating the molecular mechanisms of AT2R signaling and its interactions with AT1R. Research is also aimed at exploring the potential of AT2R as a therapeutic target. This involves the development of selective AT2R agonists and antagonists to harness its beneficial effects in cardiovascular and renal protection. Clinically, AT1R antagonists are widely used in treating hypertension and heart failure. The therapeutic potential of AT2R, however, remains under investigation. Emerging evidence suggests that AT2R agonists could offer a novel treatment strategy for cardiovascular and renal diseases, possibly providing additional benefits beyond AT1R blockade^8^. Additionally, AT2R is a potential target in glioblastoma (GBM), which suggests exploring new drug targeting AT2R can also assist cancer treatment^9^.

### AutoDock Vina

Molecular docking is a computational technique that predicts the preferred orientation of one molecule to a second when they bind to each other to form a stable complex. This method is instrumental in drug discovery, allowing researchers to model the interaction between a small molecule (often a potential drug) and a protein at the atomic level. By simulating these interactions, docking can provide insights into the binding affinities and modes of action of drug candidates, significantly aiding in the prediction of their therapeutic efficacy^10^.

AutoDock Vina is a widely-used software tool in the field of molecular docking. Notable for its improved accuracy and speed, AutoDock Vina is an evolution of AutoDock, offering faster performance by leveraging multi-threading capabilities. The software employs a sophisticated scoring function that incorporates terms for hydrophobic interactions, hydrogen bonding, electrostatics, and entropy, to estimate the binding energy of the ligand-protein complex^11^.

While molecular docking provides a static approach to predict ligand binding poses, molecular dynamics (MD) simulations offer a dynamic perspective, accounting for the full range of molecular motions over time. However, MD simulations are computationally demanding, often requiring high-performance computing to simulate the physical movements of atoms within a molecular system over time^12^. In contrast, molecular docking, particularly with AutoDock Vina, can be effectively conducted on standard personal computers. This accessibility enables researchers without access to high-powered computing facilities to still make significant contributions to understanding protein-drug interactions and to progress in drug discovery. In this paper, we use AutoDock vina to predict possible mechanism between AT2R and sparsentan.

### Sparsentan

Sparsentan (FILSPARI™) is an oral, dual-action antagonist developed by Travere Therapeutics, targeting both endothelin and angiotensin II receptors. In February 2023, sparsentan was approved to lower proteinuria in adult patients with primary immunoglobulin A (IgA) nephropathy who are facing an elevated risk of rapid disease advancement^13^.

Drawing on data from DrugBank and the FDA, sparsentan exhibits high affinity for both the endothelin type A receptor (ETAR) and the AT1R, displaying a selectivity that is greater than 500-fold for these receptors over the endothelin type B receptor (ETBR) and the AT2R. This could suggest that the primary target of sparsentan is AT1R. Intriguingly, as of March 20, 2024, the DrugBank database^14^ continues to list AT2R as a target of sparsentan, and our verification efforts show that sparsentan is not mentioned in the list of drugs targeting AT1R. This suggests a possible discrepancy in the database. Additionally, there is a notable deficiency in structural data regarding the interactions between sparsentan and the AT2R or AT1R within existing structural databases.

Based on the available data, we cannot conclusively determine that sparsentan exclusively targets AT1R. It remains unclear whether sparsentan can also target AT2R, or if there are other mechanisms of action yet to be discovered. Through this study, we aim to provide a reference that may help to bridge this gap in knowledge. Our research try to give a detailed structural interaction between sparsentan and AT2R could not only broad the understanding of sparsentan’s mechanism but also enhance the collective knowledge of therapeutics that target angiotensin II receptors.

## Results

### Parameters Optimization

In the molecular docking process, various parameters like exhaustiveness, num_modes, energy_range, and random seed can affect the outcome of the docking simulation. Of these, exhaustiveness and the random seed are two parameters that can significantly impact the results. In this study, we systematically investigated these parameters to identify the combination that yields the highest binding affinity. The results of this optimization are presented in Table.1. An exhaustiveness value of 140 and a random number seed of 312 were determined to produce the most favorable docking outcome, with the highest binding affinity recorded at -10.2 kcal/mol.

**Table 1.**
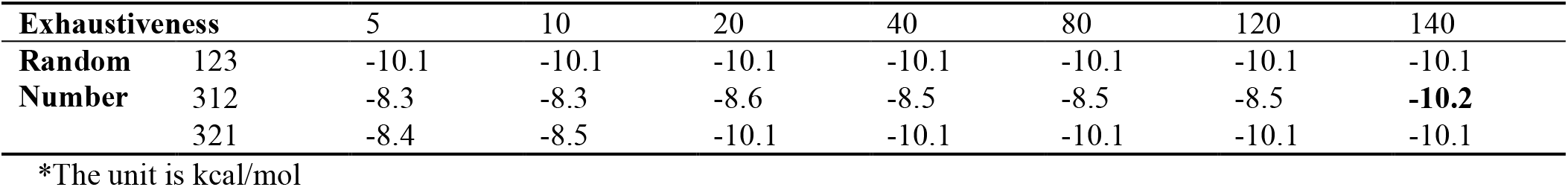
Parameters optimization for exhaustiveness and random number.

| Exhaustiveness |  | 5 | 10 | 20 | 40 | 80 | 120 | 140 |
| --- | --- | --- | --- | --- | --- | --- | --- | --- |
| Random Number | 123 | -10.1 | -10.1 | -10.1 | -10.1 | -10.1 | -10.1 | -10.1 |
|  | 312 | -8.3 | -8.3 | -8.6 | -8.5 | -8.5 | -8.5 | **-10.2** |
|  | 321 | -8.4 | -8.5 | -10.1 | -10.1 | -10.1 | -10.1 | -10.1 |

\*The unit is kcal/mol

### Overall Results

The overall results typically indicate the binding location of the drug within the protein. As depicted in Figure.1, sparsentan is fully encompassed by the AT2R, with its structure nestled among several alpha-helices.

**Figure 1.**
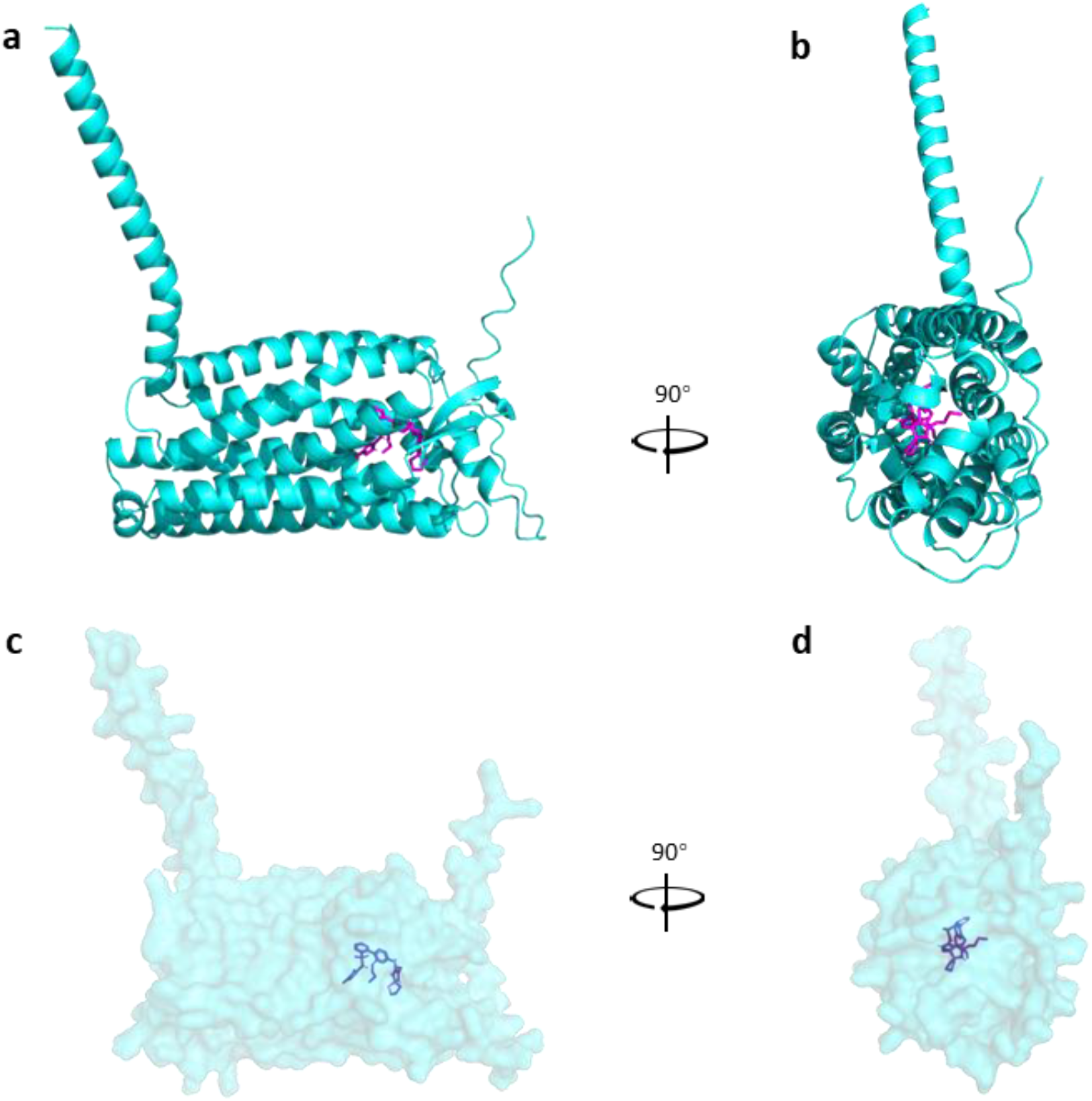
Overall results of docking. AT2R is colored in cyans and sparsentan is colored in magentas.

### Hydrogen Bond Interaction Analysis by PyMol

To analyze potential hydrogen bonds between AT2R and sparsentan, we employed the ‘find polar contacts to other atoms in object’ function in PyMol^15^. This revealed that an oxygen atom within sparsentan forms two hydrogen bonds with amino acids in the protein as shown in Figure.2. The first bond is between the oxygen atom of sparsentan and the hydrogen atom on the peptide bond of Methionine 197. The second bond occurs between the same oxygen atom in sparsentan and the hydrogen atom located at the terminal end of the side chain in Arginine 182.

**Figure 2.**
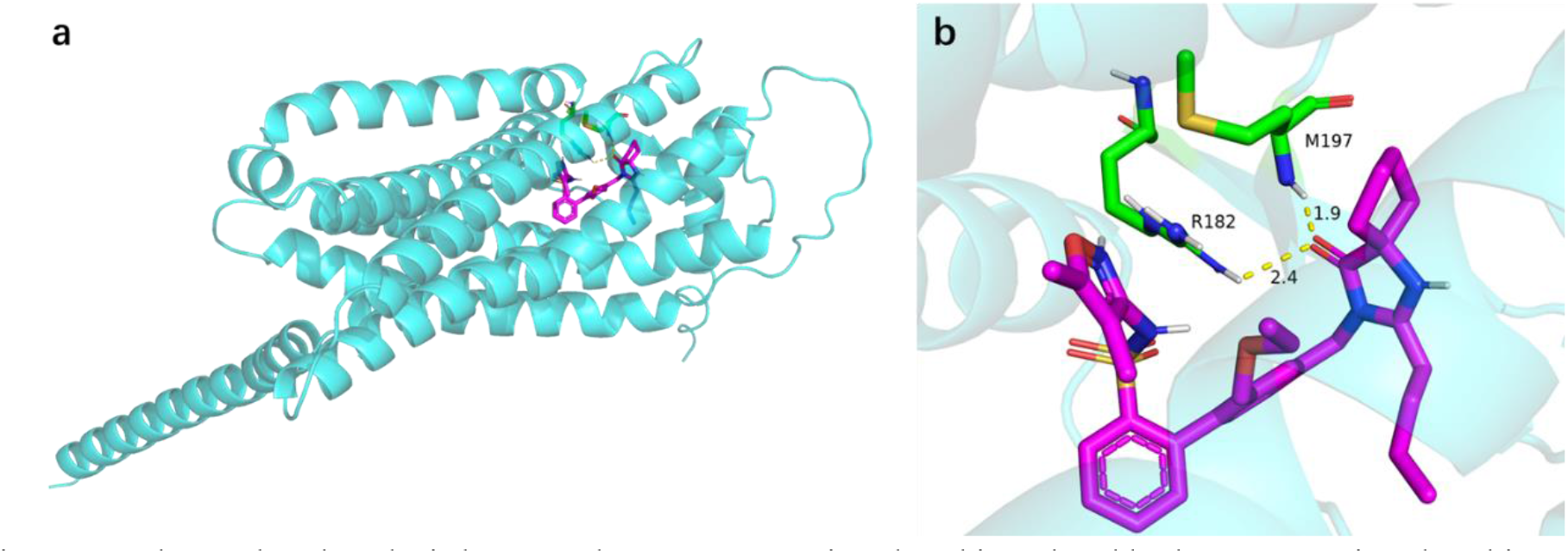
Hrdrogen bond analysis by PyMol. Oxygen atom is colored in red and hydrogen atom is colored in grey.

### Protein-Ligand Interaction Profile

The Protein-Ligand Interaction Profiler^16^, an online tool for analyzing docking results, was used to assess four types of interactions. The outcomes are illustrated in Figure.3, and the detailed results are cataloged in Table.2. It is noteworthy that the count of amino acids identified to form hydrogen bonds with sparsentan is higher by one when compared to the analysis conducted in PyMol. This discrepancy could be attributed to variations in the computational algorithms used by the respective analysis platforms.

**Figure 3.**
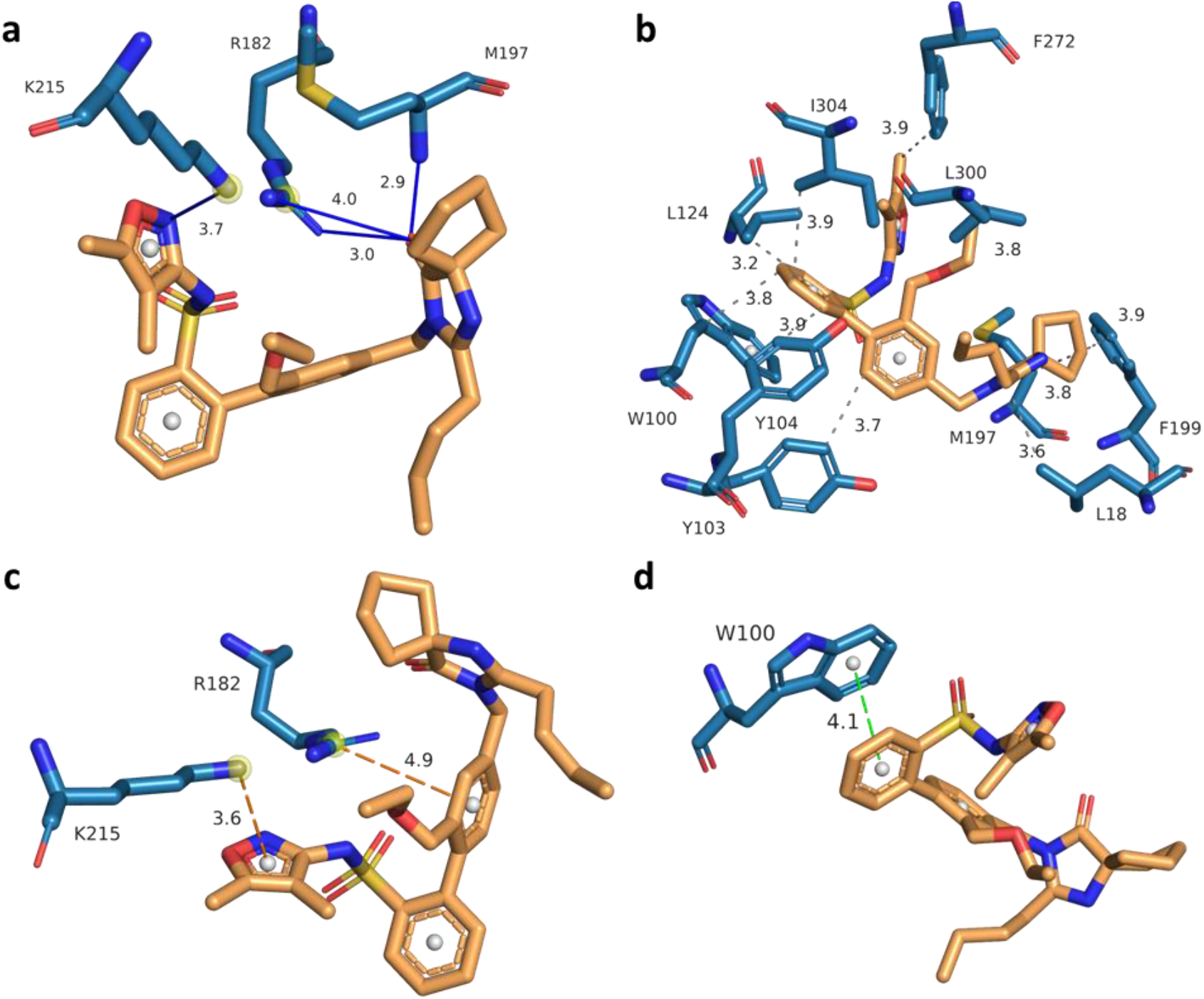
Four kinds of protein-ligand interaction. **a**. Hrdrogen Bonds. **b**. Hydrophobic interactions. **c**. π-cation interactions. **d**. π-stacking. Interaction distance is labeled with different color.

**Table 2.** Interaction profile.

|  | Index | Residue | AA | Distance<br>H-A | Distance<br>D-A | Donor<br>Angle | Protein<br>Donar? | Side<br>Chain? | Donor<br>Atom | Acceptor<br>Atom |
| --- | --- | --- | --- | --- | --- | --- | --- | --- | --- | --- |
| **Hydrogen**<br>**Bonds** | 1 | 182 | Arg | 3.59 | 4.05 | 109.33 | + | + | 1797[Ng+] | 33[O2] |
|  | 2 | 182 | Arg | 2.43 | 3.05 | 117.87 | + | + | 1796[Ng+] | 33[O2] |
|  | 3 | 197 | Met | 1.88 | 2.85 | 157.61 | + | - | 1935[Nam] | 33[O2] |
|  | 4 | 215 | Lys | 3.09 | 3.75 | 123.33 | + | + | 2107[N3+] | 19[Nox] |
|  | Index | Residue | AA | Distance | Ligand<br>Atom | Protein<br>Atom |  |  |  |  |
| **Hydrophobic**<br>**Interactions** | 1 | 18 | Leu | 3.59 | 42 | 205 |  |  |  |  |
|  | 2 | 100 | Trp | 3.78 | 10 | 963 |  |  |  |  |
|  | 3 | 103 | Tyr | 3.69 | 6 | 997 |  |  |  |  |
|  | 4 | 104 | Tyr | 3.86 | 12 | 1012 |  |  |  |  |
|  | 5 | 124 | Leu | 3.25 | 9 | 1216 |  |  |  |  |
|  | 6 | 197 | Met | 3.78 | 41 | 1939 |  |  |  |  |
|  | 7 | 199 | Phe | 3.93 | 41 | 1959 |  |  |  |  |
|  | 8 | 272 | Phe | 3.93 | 24 | 2669 |  |  |  |  |
|  | 9 | 300 | Leu | 3.83 | 29 | 2915 |  |  |  |  |
|  | 10 | 304 | Ile | 3.89 | 11 | 2948 |  |  |  |  |
|  | Index | Residue | AA | Distance | Offset | Protein<br>Charged? | Ligand<br>Group | Ligand Atoms |  |  |
| **π-Cation**<br>**Interactions** | 1 | 182 | Arg | 4.94 | 0.63 | + | Aromatic | 1,2,3,4,5,6 |  |  |
|  | 2 | 215 | Lys | 3.56 | 1.22 | + | Aromatic | 18,19,20,22,23 |  |  |
|  | Index | Residue | AA | Distance | Angle | Offset | Stacking<br>Type | Ligand Atoms |  |  |
| **π-Stacking** | 1 | 100 | Trp | 4.10 | 10.49 | 1.38 | P | 7,8,9,10,11,12 |  |  |

## Discussion

The AT2R is a crucial component of the RAS, and its mechanisms are less understood compared to those of the AT1R. There is a significant need to develop drugs that specifically target AT2R. Sparsentan is a novel FDA-approved drug designed to antagonize angiotensin II receptors. FDA documentation suggests that it has a greater affinity for AT1R compared to AT2R, yet it is listed in DrugBank as a drug targeting AT2R. Presently, structural data elucidating the mechanisms of interaction between sparsentan and either AT1R or AT2R is lacking in databases. Consequently, it is not yet possible to definitively identify whether sparsentan exclusively targets AT2R, AT1R, or if it interacts with both receptors in its mode of action.

In light of these uncertainties, molecular docking simulations using AutoDock Vina were conducted to hypothesize potential interaction mechanisms between sparsentan and AT2R. Our findings demonstrated that sparsentan binds to AT2R with a binding affinity of -10.2 kcal/mol. The interaction analysis revealed the presence of 4 potential hydrogen bonds, 10 hydrophobic interactions, 2 π-cation interactions, and 1 π-stacking interaction. The findings suggest that sparsentan is capable of interacting with AT2R, implying that its activity is not limited to AT1R alone. This could serve as an additional reference point, supplementing existing FDA documentation and DrugBank entries.

Our study highlights the possibility of a more intricate mechanism of action for sparsentan within the body, and the observed binding affinity positions sparsentan as a promising lead compound for the development of new AT2R-targeted drugs.

## Methods

### Molecular Docking and Visualization

All software applications were installed and operated on a personal computer with a Windows 10 operating system, equipped with an Intel i5-8250U processor and 8 CPU cores.

MGLTools^17^ version win32_1.5.7 was downloaded from: https://ccsb.scripps.edu/mgltools/downloads/.

AutoDock Vina version 1_1_2_win32 was downloaded from: https://vina.scripps.edu/downloads/.

The protein structure data of AT2R specific to humans was acquired from the AlphaFold^18^ database by searching for the UniProt ID: P50052.

Sparsentan was downloaded in 3D-SDF format from DrugBank and converted to PDB format using PyMol, version 2.5.7 for Windows-x86_64.

Both the protein and the drug were pre-processed with MGLTools prior to the docking procedure.

The results from AutoDock vina and Protein-Ligand Interaction Profile were visualized and analyzed using PyMol.

The protein-ligand profile was performed on Protein-Ligand Interaction Profiler. The docking results in PDB format was uploaded to the website for analysis.

All images were generated using PyMol.

### Docking Parameters Optimization

Exausitiveness and random number were set to various values in Table.1 during the docking simulation. The highest affinity result was selected to determine the optimal pair of parameters. Additional parameter details are provided in Table.3.

**Table 3.** Other parameters in molecular docking.

| Parameter | Value | Parameter | Value | Parameter | Value |
| --- | --- | --- | --- | --- | --- |
| center_x | -13.692 | size_x | 74.0 | num_modes | 20 |
| center_y | 23.115 | size_y | 100.0 | energy_range | 3 |
| center_z | 2.501 | size_z | 96.0 | cpu | 8 |

## Data Availability

The complete set of source data, including protein and drug files in PDB format, as well as docking config and interaction analysis, can be accessed at: https://github.com/Peneapple/sparsentan.

## Acknowledgements

The author declare no specific acknowledgements.

## Author Contributions

Peter X. Geng proposed the idea, researched data for this article, contributed substantially to analysis and discussion of the results, wrote the article and reviewed the manuscript before submission.

